# Porter 5: fast, state-of-the-art ab initio prediction of protein secondary structure in 3 and 8 classes

**DOI:** 10.1101/289033

**Authors:** Mirko Torrisi, Manaz Kaleel, Gianluca Pollastri

## Abstract

**Motivation:** Although secondary structure predictors have been developed for decades, current ab initio methods have still some way to go to reach their theoretical limits. Moreover, the continuous effort towards harnessing ever expanding data sets and more sophisticated, deeper Machine Learning techniques, has not come to an end.

**Results:** Here we present Porter 5, the latest release of one of the best performing ab initio secondary structure predictors. Version 5 achieves 84% accuracy (84% SOV) when tested on 3 classes, and 73% accuracy (77% SOV) on 8 classes, on a large independent set, significantly outperforming all the most recent ab initio predictors we have tested.

**Availability:** The web and standalone versions of Porter5 are available at http://distilldeep.ucd.ie/porter/.

## 1 Introduction

The prediction of protein Secondary Structure (SS) has been a central topic of research in Bioinformatics for many decades [1]. In spite of this, even the most recent and sophisticated ab initio SS predictors are not able to reach the theoretical limit of three-state prediction accuracy (88-90%), while only a few predictors are currently able to generate eight-state SS predictions.

## 2 Approach

Similarly to its previous versions [2], Porter 5 is based on ensembles of Bidirectional Recurrent Neural Networks (BRNN). Our implementation consists of two similar cascaded stages, both of which contain a classic BRNN layer followed by a convolutional layer. We have determined the hyperparameters of the system through extensive testing in five-fold cross-validation. The resulting optimal hyperparameters have been used to train the final predictor on the full training set, while the results we report are obtained on a completely independent set. The novel elements of Porter 5 are the use of HHblits [3] in its pipeline alongside PSI-BLAST [4], a more informative encoding of the inputs, larger models, deeper training and the reliance on a larger training set than in the previous versions.

## 3 Methods

### 3.1 Datasets

The selection and preparation of datasets to adopt has a central role in any machine learning method [5]. To build the training set we started from the Protein Data Bank (PDB) [6] from December 11, 2014 and redundancy-reduced it at a 25% sequence identity threshold. For testing purposes we started from the PDB released on June 14, 2017, redundancy-reduced it at a 25% sequence identity threshold against the training set, and then against itself to remove internal redundancy. Finally, we removed all proteins with at least 10 undetermined amino acids (AA) from both datasets. The training set contains 15,753 proteins and the test set 3,154 proteins, among the largest ever used to either train or test a SS predictor to the best of our knowledge.

### 3.2 Evolutionary information

The second key aspect of any SS predictor developed since the early 90’s is harnessing evolutionary information [7] in the form of profiles or position-specific substitution matrices extracted from multiple aligned sequences. PSI-BLAST [8] has been widely used to find remote homologues, and a key component of Porter since its first release [2]. More recently, HHblits [3] has distinguished itself for fast iterations and high quality results. Porter 5 relies on both algorithms, iterating them 3 times with an e-value of 0.001. PSI-BLAST is run on the Jun 3, 2016 version of UniRef90 [9]. HHblits is run on the February, 2016 version of UniProt20. Our empirical tests show similar results when a model is trained with either PSI-BLAST or HHblits, but significantly improved results when both are used, either via an ensemble or by concatenating the resulting profiles.

### 3.3 Input Encoding

Although the size of the training set is large, our experiments confirm a positive correlation between number of alignments found, and quality of the prediction [10]. We do not limit the number of alignment reported by either PSI-BLAST or HHblits, resulting in the generation of alignments with an average of ∼14,000, and ∼1,300 proteins with PSI-BLAST, and HHblits, respectively. We encode the alignments as frequency profiles. In particular we have 20 frequencies for the standard AA, 1 frequency for unknown or non-standard and one for gaps. Frequencies for AA and gaps are computed separately, that is: AA frequencies are computed ignoring gaps, while the gap frequency is equal to the total number of gaps divided by the total number of sequences aligned. Aligned sequences are weighed by the information they carry with respect to the unweighed frequency profile [11], which in our test results in a significant performance improvement. Moreover, the frequency for the AA appearing in the query sequence is artificially “clipped” to 1. This does not reduce the information of the profile, as any one of the values in the initial (unclipped) profile is equal to 1 minus the sum of the others, while encoding into the input the identity of the query sequence. We have found this simple trick to be very beneficial in our tests, leading to significant performance improvements. We also build a version of the predictor which outputs the 8 SS classes produced by the DSSP program [12]. In this case three more numbers, representing the output of the three-state Porter 5, are added to the input.

### 3.4 Model and Training

We conducted preliminary experiments to identify optimal hyperparameters using only the training set in 5-fold cross-validation. In this phase we also extensively tested many types of neural network, including convolutional networks of increasing depths and simple windowed neural networks with up to 17 hidden layers, but our two-stage BRNN/CNN model [2] performed over 1% better than any of the alternatives we tried. The hyperparameters selected for the BRNNs trained on PSI-BLAST appeared close to optimal for both the BRNN trained on HHblits, and the BRNN trained on the concatenation of the inputs created with HHblits and PSI-BLAST. Only a slight increase of the hyperparameters was needed when switching from three-state to eight-state SS. Purely based on the training set we settled on an ensemble of 7 BRNN: 3 trained on PSI-BLAST, 3 trained on HHblits, and 1 trained on both (44 concatenated inputs rather than 22). These 7 BRNN were then trained on the full training set (on either three-state, or eight-state targets). The relatively small scale of the ensemble and the modest size of the individual models (on average 39k free parameters for the 3-class networks, 58k for 8 classes) ensure high speed at prediction time without affecting the accuracy or SOV score (Table 1). The test set was only used at the final stage, to obtain unbiased performance estimates for Porter 5 and other recent SS predictors [5].

**Table 1:** Performances on the set for which SPIDER3 generates predictions, sorted by Q3 accuracy.

| Method | Q3<br>per AA | SOV'99<br>per AA | SOV_refine<br>per AA | Q3<br>per protein | SOV'99<br>per protein | SOV_refine<br>per protein |
| --- | --- | --- | --- | --- | --- | --- |
| **Porter 5** | **83.81%** | **80.41%** | **75.73%** | **84.32%** | **81.05%** | **76.45%** |
| SPIDER3 | 83.15% | 79.43% | 74.68% | 83.42% | 79.79% | 75.07% |
| Porter5 (HHblits only) | 83.06% | 79.49% | 74.71% | 83.68% | 80.26% | 75.58% |
| SSpro 5 with templates | 82.58% | 78.54% | 74.02% | 83.94% | 80.29% | 76.15% |
| PSIPRED 4.01 | 81.88% | 77.36% | 72.33% | 82.48% | 78.22% | 73.31% |
| RaptorX-Property | 81.86% | 78.08% | 72.99% | 82.57% | 78.99% | 74.03% |
| Porter 4 | 81.66% | 78.05% | 72.89% | 82.29% | 78.61% | 73.55% |
| SSpro 5 ab initio | 81.17% | 76.87% | 72.03% | 81.10% | 76.92% | 72.12% |
| DeepCNF | 81.04% | 76.74% | 71.47% | 81.16% | 76.99% | 71.7% |

## 4 Results

We tested Porter 5 against Porter 4 [13], SPIDER3 [14], SSpro 5.2 [15], PSIPRED 4.01 [16], RaptorX-Property [17] and DeepCNF-SS [18] on the test set we created, containing 3,154 proteins. However we were limited in this by the fact that SPIDER3 rejects proteins containing undetermined (X) amino acids (562 in the test set contain at least one), and that when we use the parameters required by SPIDER3, either PSI-BLAST or HHblits do not return a valid result for 129 proteins. Because of this we report performance results on two sets: one where we exclude the proteins on which we could not obtain a valid response from SPIDER3, containing 2,463 entries in total (Table 1); the full set of 3,154 proteins (Table 2) on which SPIDER3 is not assessed.

**Table 2:**
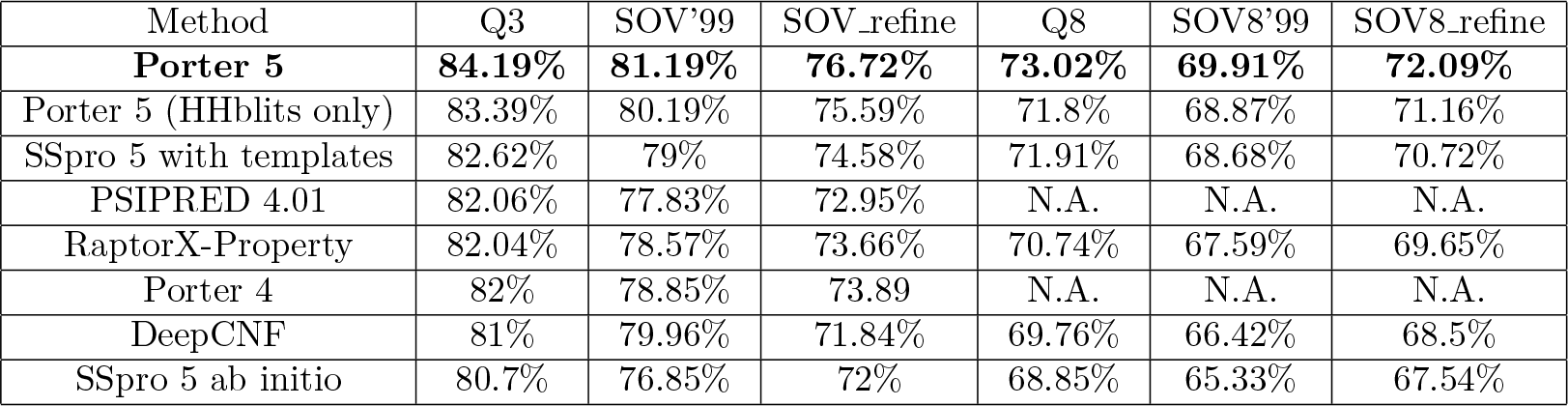
Q3/Q8 accuracy and SOV’99 and SOV_refine score per AA on the full test set.

| Method | Q3 | SOV'99 | SOV_refine | Q8 | SOV8'99 | SOV8_refine |
| --- | --- | --- | --- | --- | --- | --- |
| **Porter 5** | **84.19%** | **81.19%** | **76.72%** | **73.02%** | **69.91%** | **72.09%** |
| Porter 5 (HHblits only) | 83.39% | 80.19% | 75.59% | 71.8% | 68.87% | 71.16% |
| SSpro 5 with templates | 82.62% | 79% | 74.58% | 71.91% | 68.68% | 70.72% |
| PSIPRED 4.01 | 82.06% | 77.83% | 72.95% | N.A. | N.A. | N.A. |
| RaptorX-Property | 82.04% | 78.57% | 73.66% | 70.74% | 67.59% | 69.65% |
| Porter 4 | 82% | 78.85% | 73.89 | N.A. | N.A. | N.A. |
| DeepCNF | 81% | 79.96% | 71.84% | 69.76% | 66.42% | 68.5% |
| SSpro 5 ab initio | 80.7% | 76.85% | 72% | 68.85% | 65.33% | 67.54% |

Porter 5 is the most accurate predictor in our tests with 3-class accuracy just shy of 84% on the smaller testing set and 84.2% on the larger set, 0.7% better than SPIDER3, 1.2% and 1.6% better than SSpro with templates, and at least 2% more accurate than all the other predictors. Porter 5 is also very fast compared to the alternatives given the relatively small size of its models and the fact that it is built on in-house, heavily profiled code. For instance, once the alignments by PSI-BLAST and HHblits are present, Porter 5.0 runs 2 orders of magnitude faster than SPIDER3, although SPIDER3 is able to predict backbone angles, contact numbers, and solvent accessibility at the same time as secondary structure. To fully exploit its speed, we also assessed, and make available, a version of Porter 5 which is roughly three times faster depending on HHblits only.

It should be noted that our test set is redundancy reduced only against our training set, so our assessment of other recent SS predictors might be somewhat optimistic, as the test set may contain proteins similar to those they were trained on. Porter 5 is the best three-state and eight-state SS predictor according to all the measures observed on this set. Porter 5 with HHblits only performs better than SSpro 5 with templates, at least 1.2% better than RaptorX-Property (the only other predictor based on HHblits only), and has similar performances to SPIDER3.

We have also analysed the performance of Porter 5 separately on proteins resolved by Nuclear Magnetic Resonance (NMR) and by X-ray crystallography. NMR proteins are predicted at a significantly lower Q3 (81.6%), possibly because of their different statistics (e.g. average length and composition) or less certain determination of SS. The X-ray only section of the test set, which is roughly 90% of the total, is predicted at an average Q3 of 84.65% (Table 3).

**Table 3:** Porter 5 on NMR vs X-ray crystallography proteins

| Method | Q3 | SOV | Q8 | SOV |
| --- | --- | --- | --- | --- |
| NMR | 81.61% | 78.23% | 67.52% | 70.71% |
| **X-ray** | **84.65%** | **85.08%** | **74%** | **77.93%** |

## 5 Conclusion

We have built a state-of-the-art 3-and 8-state predictor of protein SS, Porter 5. In our tests it improves by roughly 2% accuracy on its previous version, and outperforms all the most recent predictors of secondary structure, including a template based one. Porter is freely available as a web server at http://distilldeep.ucd.ie/porter/ and also as a standalone program at the same address, alongside with all the datasets, and alignments generated.

## Acknowledgements

We acknowledge the key role of the UCD cluster, and the UCD Research IT staff. We also acknowledge the use of the SOV_refine software to calculate all SOV scores[19].

